# Harmonizing resting-state functional MRI data to increase sample size and to classify tinnitus brain connectivity

**DOI:** 10.64898/2026.11.19.767894

**Authors:** Shagun Ajmera, Fatima T. Husain

## Abstract

Tinnitus, or ‘ringing in the ears’ is a hearing disorder that affects approximately 15% of human adults globally. The neuroscientific understanding of this subjective disorder is limited at present, partly due to a lack of sufficient neuroimaging data to compensate for the heterogeneity in the disorder; neural data acquisition is often impeded by low patient counts and doubled costs against scanning ample controls. However, with the increase in data sharing practices, data scarcity can be overcome with ‘harmonization’ methods if scans from independent sites and studies are pooled successfully. To achieve data harmonization for limited data from a tinnitus neuroimaging study, we merged a modest fMRI dataset acquired in our lab (n=35) with age- and gender-matched samples from another large dataset, the Lifespan Human Connectome Project Aging (n=377). We measured functional connectivity (FC) between brain regions based on resting-state fMRI session scans, and used deep learning architecture to systematically combine individual FCs across the datasets. We assembled and trained neural network modules to purge dataset-identifying information from FC maps, and retained patterns that can characterize tinnitus-related FC. Hallmark artifactual information of the datasets was significantly reduced while reconstructing resting state FC data. Relevant relationships in brain connectivity that discern the condition of tinnitus from controls were well-preserved in the reconstructed resting state FC data. In agreement with existing literature on brain networks of tinnitus, connectivity related to default mode, ventral attention and frontoparietal control networks were identified to be crucial for distinguishing the condition, as masking these networks’ connections severely impacted downstream classification prediction on reconstructed FC. The data harmonization model was successful in merging FC matrices from independently acquired fMRI datasets, without losing out on meaningful functional connectivity patterns of interest for the neuropsychological disorder of tinnitus. Our findings demonstrate the potential of data harmonization by integrating independently acquired datasets on healthy controls to overcome neuroimaging data scarcity while preserving disorder-relevant neural signatures, advancing research on understudied neurological disorders.

## Introduction

Although MRI is an indispensable neuroimaging tool for investigating human brain functions and dysfunctions, high costs associated with image acquisition is one of the major disadvantages of the method, due to which, gathering large amounts of MRI scans becomes a tough challenge. Moreover, for studying hearing-related disorders such as tinnitus, functional MRI data collection is further hindered by scanner-related logistics (ex: bothersome scanner noise) and small number of participants in the patient cohort.

However, thanks to open data policies and large neuro consortiums, there is increasing availability of pre-acquired anonymized MRI human data. It is agreed upon that data sharing can be resourceful for advancing neuroimaging based research, especially for progressing our understanding of auditory cognitive disorders (Eckert et al., 2023). The often debilitating condition of tinnitus is one such hearing disorder that affects more than 740 million adults globally (Jarach et al., 2022). The United States Department of Veterans Affairs bears highest compensation costs annually against tinnitus, and recognizes tinnitus as the most prevalent service-connected disability (Veterans Benefits Administration, 2023). Phantom auditory perception can be bothersome to tinnitus sufferers in their day-to-day lives through psychological, cognitive, and sleep issues (Hall et al., 2018; Langguth et al., 2011; Watts et al., 2018); yet the neural bases associated with this condition remain under-studied. Despite the financial and mental toll of tinnitus, research in tinnitus remains under-funded, which is also a bottleneck for ample data collection.

When dealing with limited patient data, which is the case with tinnitus studies, a useful innovation could be to leverage other open access databases for benchmarking different brain health conditions. A few recent studies have exploited computational techniques for hyperalignment of MR-acquired data from multiple sources (Guntupalli et al., 2018; Nastase et al., 2020; Yamashita et al., 2019). These example studies demonstrate that, at the data quality front, aggregating across different data repositories can not only relieve statistical demands, but also enable recognizing and eliminating scanner- or site-related biases in the data.

A primary issue in merging different MRI datasets is that the aggregation is not straightforward, as there can be dataset-intrinsic noise in the scans, owing to differences in scanner types, acquisition parameters, and samples of subjects. Therefore data-’harmonization’ is a crucial precursor task to mining pooled MRI-based data for neural patterns of interest.

In a recent study by An et al. (2022), the authors combined anatomical T1 scans across three large-scale databases for Alzheimer’s disease. They harmonized the brain volumes using a model adapted from Moyer et al. (2020) which involved a conditional variational autoencoder in conjunction with an adversarial discriminator. They expanded the model further by adding a goal-directed module that predicted clinical diagnosis and behavioral scores; the augmented model significantly improved downstream prediction performance on the harmonized brain volume data.

In the present study, we implemented and tested goal-directed data harmonization on functional connectivity (FC) data. We utilized the publicly available database of Lifespan Human Connectome Project Aging (HCA database) (Bookheimer et al., 2019). We pooled together FC datasets derived from the fMRI runs in HCA with our in-house data of tinnitus patients and controls (Lab database). We tested whether the two independently acquired datasets could be harmonized, honoring our broader goal of characterizing tinnitus-related brain connectivity. Our harmonization framework not only removed dataset differences from the original data but also reliably discerned meaningful cognitive interactions in the brain for the hearing disorder of tinnitus.

### Harmonization of independent fMRI functional connectivity data

Resting state FC vectors were obtained from both the fMRI databases, and used in model training and testing steps of the harmonization process.

#### Participant matching

In the Lab dataset, multiple resting state fMRI sessions were acquired across participants in tinnitus (n=15) and control (n=20 conditions (43.1±10.8 years, 43% male) (Table 1). All Lab participants had normal hearing sensitivity as measured using comprehensive audiological assessments. From the HCA dataset, 377 subjects matched in age (+/-1 year) and gender with at least one of the Lab participants, yielding approximately 21 matched subjects per Lab participant. Hearing health profiles of HCA subjects were not available; all HCA subjects were identified as healthy individuals in the original study (Bookheimer et al., 2019), so we placed our HCA cohort in the controls category (Table 1). FC matrices were obtained for the available resting state runs by calculating Pearson’s correlation coefficient between 100 brain regions of interest (ROIs) using Schaefer atlas (Schaefer et al., 2018). Thus, a datapoint for the model was an FC vector with 4950 edges or connectivity measures.

**Table 1:** FC data used in the model was estimated on resting state sessions from Lab participants, and age-, gender-matched HCA participants.

| **Dataset** | **Category** | **# subjects** | **# rs-sessions** |
|---|---|---|---|
| Lab | Tinnitus | 15 | 56 |
|  | Controls | 20 | 76 |
| HCA | Controls | 377 | 1504 |

#### Harmonization framework

The architecture used for resting state FC harmonization is depicted in Figure 1. At the crux of the model was a variational autoencoder (VAE) which consisted of (1) an encoder whose input was an FC vector, or ‘original’ FC, (2) a decoder whose output was the corresponding decoded or ‘reconstructed’ FC vector, having same dimensions as the input, and (3) a bottleneck layer in between, which represented mean and standard deviation for the encoding projected in latent space. The reconstruction loss between estimated FC and its corresponding original FC was measured using mean squared error (MSE).

**Figure 1:**
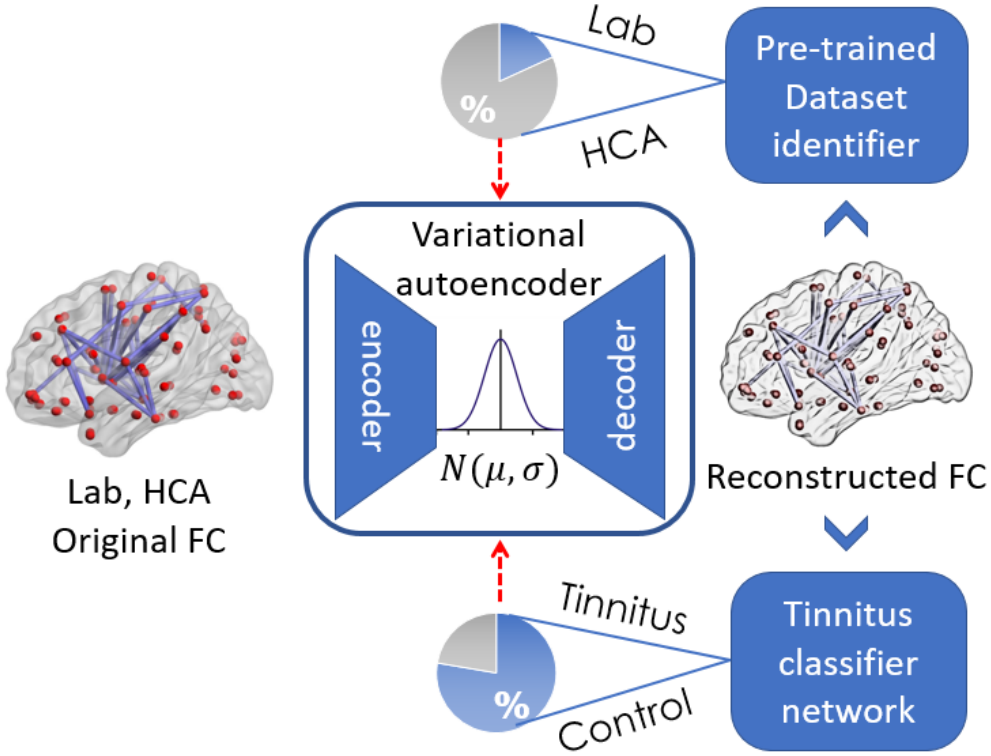
FC harmonization framework. Dashed red lines depict additional error terms used for updating VAE weights.

A logistic regression model was pre-trained on original FC to predict whether a given FC belonged to HCA dataset or Lab dataset. For harmonization training, the pre-trained model evaluated the source dataset for reconstructed FCs, and the predicted probability was used in the VAE loss term.

A feed-forward network was used to classify the reconstructed FC into tinnitus or control, and was trained concurrently with the VAE. Reconstructed FC were input to the classifier network, and the binary cross-entropy error in tinnitus/control prediction was used not only to train the classifier network, but also to tune the weights for encoder and decoder of the VAE.

During the harmonization model training phase, gradients for the VAE were driven by (1) MSE measured between the reconstructed FC and the original FC, and (2) the absolute difference between chance prediction of 0.5 and predicted probability of dataset source (|0.5 − *p*_*dataset*_|) for the reconstruction. Gradients for the encoder and decoder weights were additionally driven by the tinnitus classification error.

#### Dataset identification

A multivariate logistic regression-based classifier was trained to identify the source dataset from FC, with the dependent variable encoded as 1 for HCA and 0 for Lab. The model was trained over multiple iterations by bootstrapping samples for train and test data. Only the (original) FC data from Lab controls and their matched HCA subjects were used in training/testing the dataset identifier. Tinnitus participants’ FC was not used so as to ensure that the classifier does not learn to predict a data source by simply exploiting disorder-related patterns in FC. To train the dataset identifier, we first divided the Lab-control FC data into train and test datasets (3:1). We then added one randomly chosen FC from HCA corresponding to each train and test Lab control based on age and gender match. We performed model training over 20 repetitions by shuffling the Lab controls between train and test data, and including matched HCA controls in the same manner each time.

The average model (model coefficients averaged across 20 repetitions) correctly classified 100% of all Lab controls FC (probability prediction < 0.5) and 93.35% of all matched HCA FC (probability prediction > 0.5) (Figure 2 left-side violin plots). The averaged model was used as a pre-trained dataset identifier during the learning phase of the harmonization model (Figure 1). While training the harmonization model, the pre-trained dataset identifier evaluated data source probability for the reconstructed FC, and the evaluation was used to inform the VAE about presence of source-related components in the reconstruction. Specifically, the loss function of VAE included a ‘dataset identifiability’ measure, |0.5 − *p*_*dataset*_|, where *p*_*dataset*_ was the predicted probability that the FC belongs to HCA dataset.

**Figure 2:**
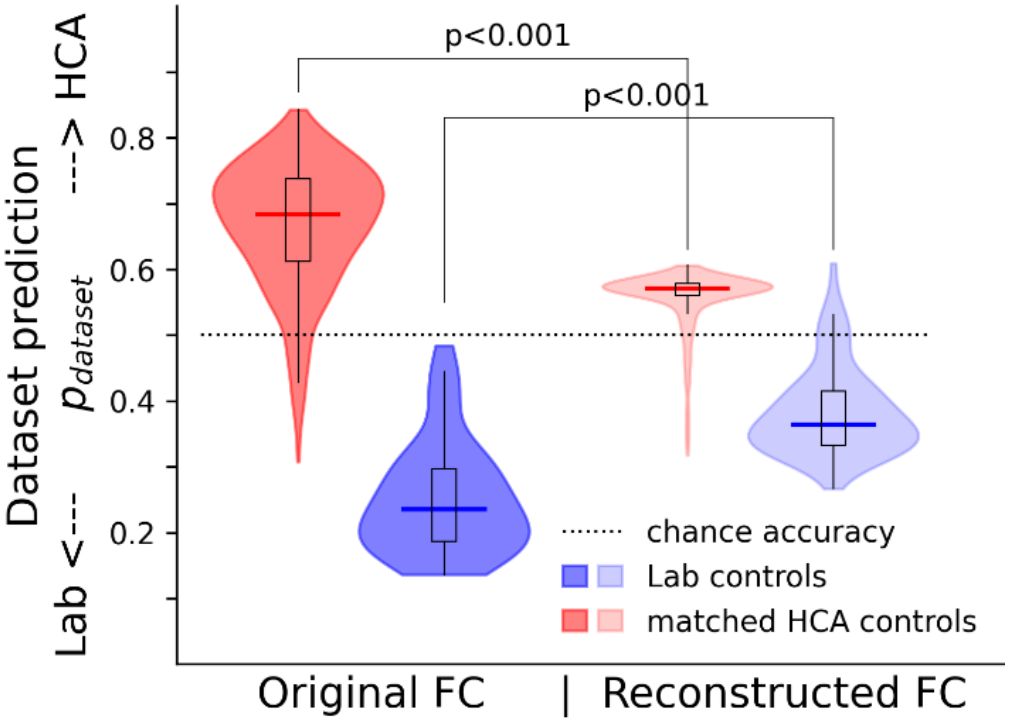
Dataset prediction for Lab and HCA controls before (left) and after (right) harmonization.

After the harmonization model training was completed, we compared dataset probability predictions on FC. As an effect of the added term on dataset identifiability, minimization of VAE loss function drove the dataset prediction for reconstructed (harmonized) FC to be significantly less accurate (closer to chance) as compared to the original FC (Figure 2 right-side violin plots) for both datasets. Thus, dataset-specific information was removed to a significant extent in the harmonized FC.

#### Autoencoder performance

The VAE unit in the harmonization model mapped an original FC vector onto a lower dimensional probability distribution, and built back the FC from a sample point in the distribution (Figure 1). The VAE was trained for 500 epochs, and over 20 repetitions by bootstrapping samples for train and test data. At the beginning of each training repetition, we first shuffled the Lab tinnitus FC data and split into 3:1 for train and test respectively. For train and test control data, we chose twice as many FCs by sampling one FC datapoint from a Lab-control and one from an HCA subject who matched in age and gender with each of the corresponding train/test tinnitus participants.

Figure 3 shows the VAE loss curves during training epochs, for the reconstruction error (MSE) and the dataset identifiability of reconstructed FC (|0.5 − *p*_*dataset*_|), and across train (Figure 3 left) and test (Figure 3 right) FC data. Loss curves for the 20 repetitions are represented in Figure 3 with overlaid translucent lines; white lines show the average error across repetitions.

**Figure 3:**
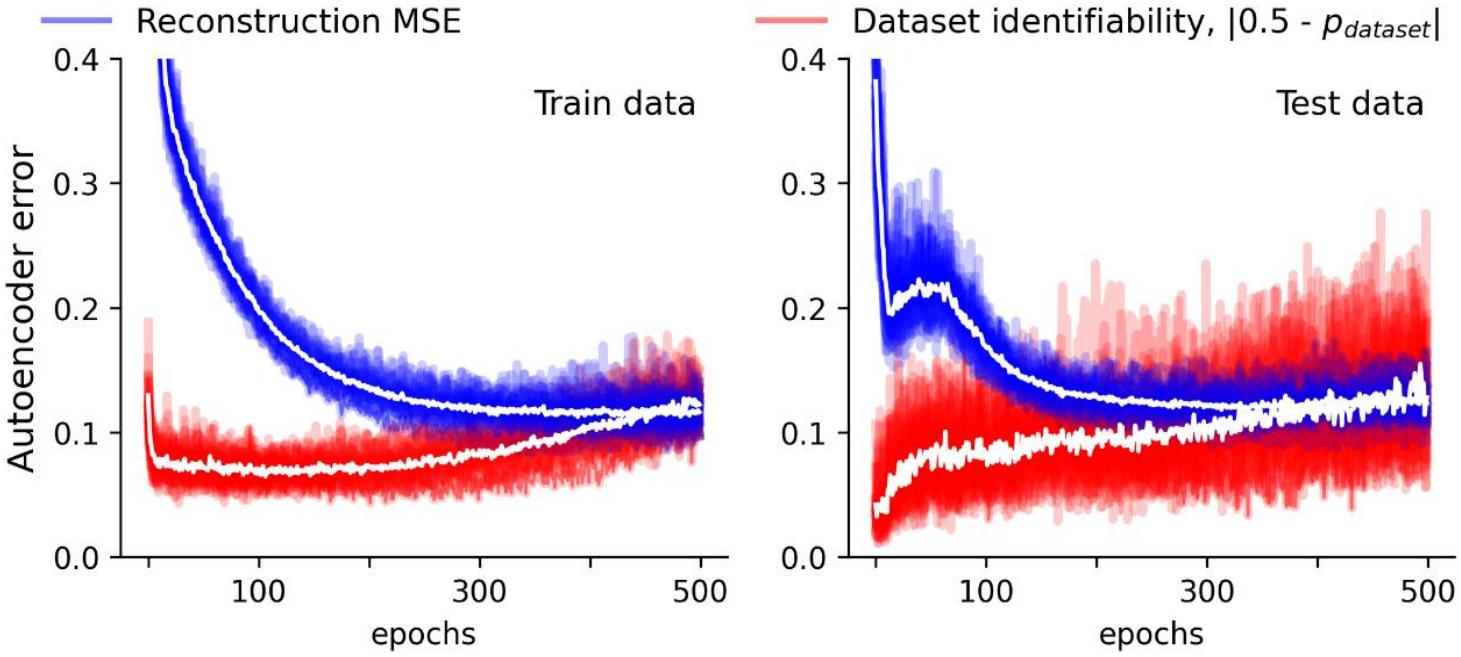
VAE training. Reconstruction error & dataset identifiability of reconstructed FC across training epochs

The quality of FC reconstruction improved with more training epochs, as evidenced by the decreasing MSE error in both the train and test data as well as across the independent training repetitions.

Dataset identifiability of reconstructed FC reduced quickly in the early epochs for train data. After the training, dataset prediction performance on reconstructed FC was desirably significantly reduced compared to original FC for controls in both the datasets (p<0.001; Figure 2) as well as for Lab tinnitus group (p<0.001). We speculate that the source dataset information in FC can be further minimized or completely removed during reconstruction by using a powerful, more complex model for dataset identification, instead of a monotonic logistic regression-based model.

### Goal-directed training of harmonization model to identify tinnitus-related FC

We trained our harmonization model to align individual FCs with reduced source-related information. We also trained the model to preserve signatures of cognitive importance in FC, such as the characteristic brain interactions indicative of tinnitus.

#### Tinnitus classification

Concurrent with the VAE training, we trained a feedforward fully-connected artificial neural network for tinnitus classification. The prediction of VAE on its train and test data (i.e. the reconstructed FCs) at each epoch were used as train and test data respectively in tinnitus classifier training. The output layer had 2 units and was followed by a softmax operation to yield probabilities corresponding to tinnitus/control labels. Binary cross entropy loss function was used to quantify error between predicted class label probability and the true label (corresponding to original FC). Tinnitus and control categories were coded using one-hot encoding.

In our harmonization model framework, the classifier error term was included in the loss function of VAE, such that the gradient estimates for encoder and decoder modules were dependent on tinnitus classifier prediction (in addition to reconstruction loss and dataset identifiability; Figure 1). This way, VAE module was trained in unison with the tinnitus classifier; Supplementary Figure 1 shows the classification error of the reconstructed FC during training phase.

After training phase completion, FC reconstructed by the goal-directed harmonization model showed more accurate category predictions (median predicted probability increased for tinnitus category, and decreased for control; Figure 4 lighter plots). Tinnitus classifier, which was trained on reconstructed FC, showed poorer category predictions on the original FC (p<0.001; Figure 4 darker plots). This suggests that the harmonized version of FC retained or enhanced statistical patterns among tinnitus-identifying functional connections, which may not manifest as evidently in the unharmonized counterpart.

**Figure 4:**
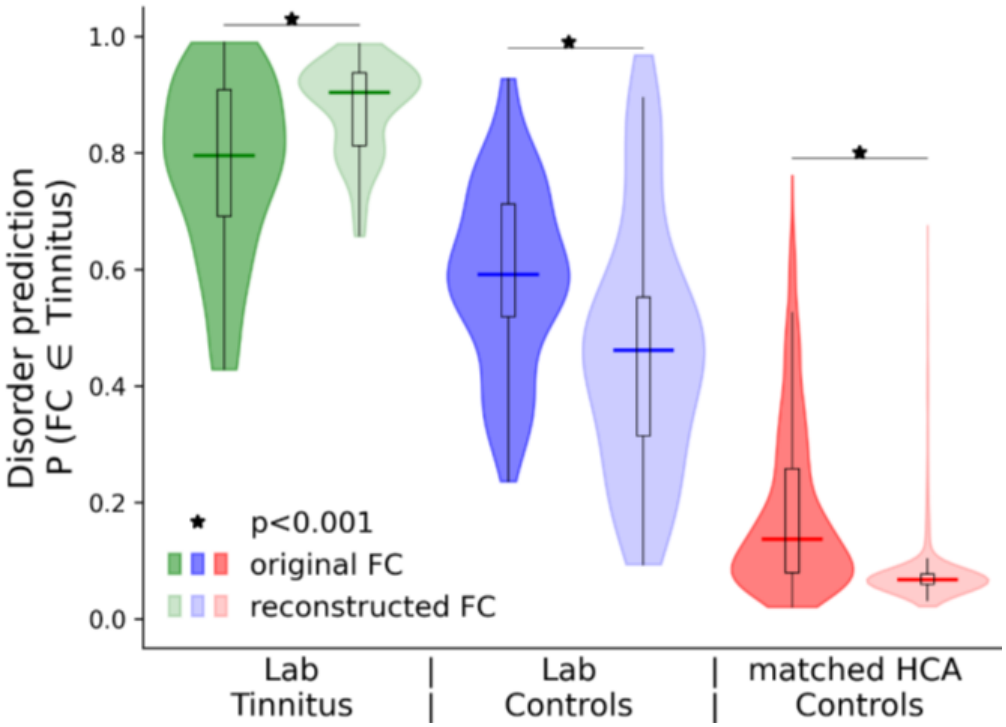
FC Tinnitus classification before (original) and after (reconstructed) goal-directed harmonization.

#### Tinnitus brain network connectivity

Given the promising performance of FC data reconstruction and its tinnitus classification, we investigated whether tinnitus detection in the reconstructed FC was driven by meaningful network connections, or by artifactual patterns that could have been induced in the data as a result of VAE decoder transformations from a lower dimensionality to actual FC space. To test the same, we evaluated the role of intrinsic network connectivity on tinnitus classification performance for the harmonized FC. We grouped the ROIs into their respective functional networks as per Yeo atlas (Thomas Yeo et al., 2011). We evaluated tinnitus classification predictions by disregarding the FC edges for one functional network at a time and using only the remaining edges of other networks in the brain. This was accomplished by selectively enforcing classifier input layer weights to zero for those FC edges that connected any ROI belonging to a chosen network, thereby completely removing both within- and between-network connectivity for the network. We call this ‘masking’ where input from a particular network (edges) is made unavailable to the classification model.

Figure 5 shows tinnitus classifier prediction on the harmonized Lab tinnitus FC data, when all the edges were intact (full FC; dark green violin plot) and when one of the functional networks’ connectivity was masked (light green violin plots). Masking five of the eight functional networks showed a significant drop in probabilities predicted for the tinnitus category (p<0.001). Interestingly, when default mode network (DMN) connectivity was occluded from harmonized FC, classifier predictions dropped most significantly (Figure 5 second plot). Considering the contribution of individual networks, DMN was the most critical; masking DMN connections alone lead to the greatest reduction in classification accuracy (∼70% decrease in median probability compared to full FC).

**Figure 5:**
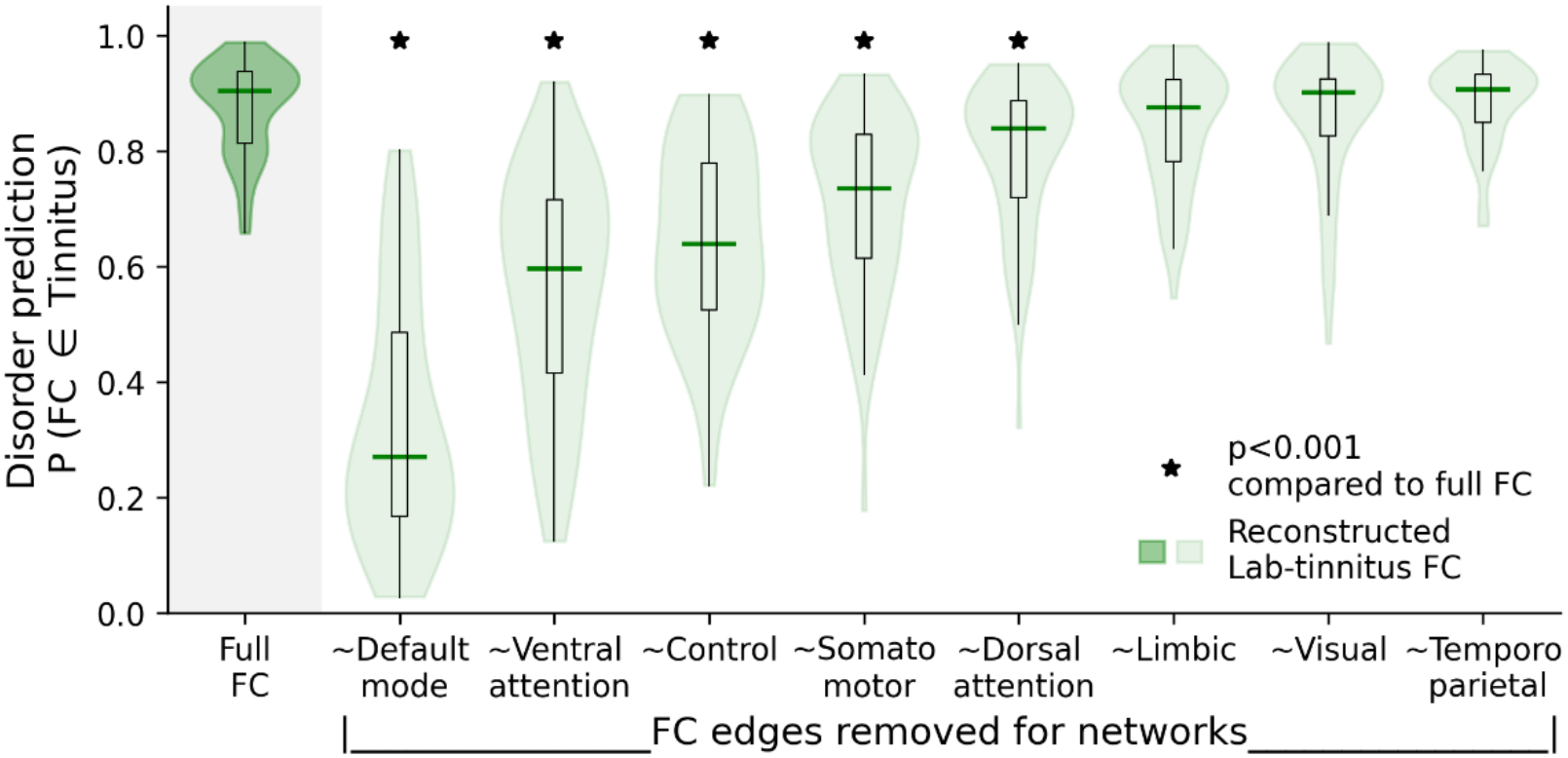
Tinnitus classifier prediction on harmonized Lab tinnitus FC data, with all FC edges intact (first plot on the left) and with FC edges of functional network ROIs removed (eight plots on the right). Functional network membership for ROIs was determined with Yeo et. al (2011) parcellation.

Complementary masking the network edges, we also performed classification by considering edges from only one network at a time. The results corroborated the prominent effect of DMN-related connectivity in tinnitus identification (Supplementary Figure 2). Classifying using DMN FC alone yielded significantly better mean probability prediction for the tinnitus cohort than did any other network.

## Discussion

In this study, we developed a goal-directed, deep learning-based harmonization framework to integrate resting-state functional connectivity data from independently acquired fMRI datasets while preserving neural signatures relevant to tinnitus. Our approach not only reduced dataset-specific biases but also enabled the integration of a small tinnitus cohort with a larger dataset of healthy controls, facilitating the identification of disorder-related connectivity patterns. Using this framework, we identified the prominent contribution of default mode network connectivity, alongside attention and cognitive control networks, in distinguishing individuals with tinnitus from controls. Below, we discuss the algorithmic advances underlying FC harmonization, their contributions to understanding the neural bases of tinnitus, and the broader implications and limitations of this approach.

### Algorithmic advances in FC harmonization

Our harmonization framework integrated resting-state FC data from a small tinnitus cohort with a large, independently acquired dataset of healthy controls. This approach increased the available sample size, introduced greater variability in the control group, and facilitated the identification of tinnitus-related connectivity patterns while reducing dataset-specific biases. Successful goal-directed FC harmonization was achieved through the synergistic integration of three key algorithmic components.

First, a logistic regression model identified statistical patterns in FC data associated with dataset-specific differences. Using this pre-trained dataset classifier as a critic, the harmonization framework reduced, although did not completely eliminate, dataset-specific information during FC reconstruction. We speculate that pre-training a more complex deep neural network-based classifier could capture additional patterns of dataset-related variability and further minimize source identifiability in the reconstructed FC.

Second, autoencoder training progressively improved the quality of reconstructed FC, as evidenced by declining reconstruction error across training epochs and repeated training iterations. Variational autoencoders map input data into a lower-dimensional latent space, with a bottleneck layer representing the parameters of a latent probability distribution, such as a Gaussian distribution (Kingma & Welling, 2019). A key component of conventional VAE training is Kullback–Leibler divergence (KL error), which regularizes the latent representation. Although our current model was trained by minimizing reconstruction error, dataset identifiability, and tinnitus classification error, incorporating KL divergence could further improve its statistical properties (Sohn et al., 2015).

Third, the tinnitus classifier guided the harmonization model to preserve disorder-relevant brain connectivity patterns in the reconstructed FC. Classification error was propagated back to the encoder and decoder modules, thereby directing FC reconstruction toward retaining information relevant to distinguishing tinnitus from controls. This goal-directed component was particularly important because it enabled the model to reduce dataset-specific variability without compromising the functional connectivity signatures relevant to the neurological condition.

### Cognitive networks of tinnitus

Harmonized FC was successfully reconstructed with the goal-directed framework. Connectivity associated with the default mode network (DMN) was identified to be most crucial in differentiating tinnitus from controls; this result corroborates previously reported markers and correlates of tinnitus in spontaneous neural connectivity of DMN among individuals with chronic tinnitus and varying degrees of severity and habituation (Rosemann & Rauschecker, 2023; Schmidt et al., 2013, 2017).

Interestingly, connectivity of salience/ventral attention network, (frontoparietal) control network, and dorsal attention network also emerged significant in tinnitus classification, as the classifier performance reduced significantly in absence of these network connections. This speaks to alterations in the interplay of attention processing and cognitive control in tinnitus (Roberts et al., 2013) because greater modifications in the dynamics of these networks are known to be linked to more bothersome tinnitus (Shahsavarani et al., 2021; Wineland et al., 2012).

Lastly, connectivity of somatomotor network also significantly affected tinnitus classification performance, albeit to a much smaller extent than the other networks discussed above. Auditory cortex and parts of somatosensory system, which are implicated in tinnitus perception (Sabados et al., 2025; Vanneste et al., 2011), were also included in this network as per the brain parcellation used for this analysis. Therefore it was an expected finding that the connectivity patterns associated with ROIs under this network contributed in tinnitus detection.

Brain imaging combined with powerful analyses and ample data can aid in extracting the idiosyncrasies of networks interactions reliably. The classifier developed in this study utilized harmonized data and shed light on brain-wide alterations in tinnitus among connections involving distinct large-scale functional networks. This perspective is important in accelerating tinnitus research as previous studies have largely tended to overlook complex network reconfigurations in tinnitus (Eggermont, 2015), which indeed manifest at a whole-brain level rather than within anatomically or functionally isolated brain regions. Evidently, harmonization can boost reliability when studying interconnectedness among cognitive networks of significance in behavioral disorders disadvantaged by smaller patient population or limited neuroimaging data.

### Broader potential

Neuroimaging studies of under-researched disorders are often limited by lower recruitment in patients, and high costs to conduct experimental fMRI sessions on patients and contemporary controls. Supplementing the data, with similar open access fMRI datasets on control groups and/or patient populations, is a viable solution, which can be challenging because differences in scan configurations, recruitment strategies, and processing pipelines can induce site-specific variability and noise or biases in the datasets. Our harmonization framework was useful in addressing some of the irrelevant variance, and extracting more robust neural patterns of relevance for the disorder. Our use case in tinnitus motivates the potential application of fMRI FC harmonization for characterizing altered cognitive networks among the broad class of brain disorders with behavioral abnormalities.

Resting state fMRI functional connectivity data from independently acquired databases were harmonized with the combined arrangement of a pre-trained dataset identifier, an autoencoder, and a tinnitus classifier. Systematic goal-directed training of the harmonization model helped distinguish individuals with tinnitus from controls across datasets, by removing dataset-related noise, and accentuating the informative patterns in intrinsic brain interactions that signify presence or absence of the disorder. Our results testify the promising use of FC harmonization for consolidating independently acquired fMRI datasets. Moreover, the FC harmonization scheme may be particularly beneficial for studying common connectivity patterns across the smaller subtypes of a heterogeneous condition (example: autism spectrum disorder), or disorders with relatively small patient population (insufficient neuroimaging data; example: Tourette syndrome).

### Limitations

The present model is not agnostic to the number of datasets to be harmonized. Training from scratch may be required for pooling newer datasets in future, in order to recognize and remove any irrelevant features that are specific to the newer data source(s). While our study illustrates data harmonization between different cohort sizes (varying number of datapoints available per group and per site), future work should generalize the framework by adapting the model to multiple datasets (not just two). Comparison with other harmonization tools like ComBat (Fortin et al., 2017, 2018; Johnson et al., 2007) is also necessary to realize the strengths and weaknesses of our deep-learning based FC harmonization model. Additionally, an adversarial neural network such as GAN (Goodfellow et al., 2014) can be trained in parallel with the VAE to possibly achieve a more persistent decline in dataset identifiability over the training epochs. Such improvements in model architecture remain to be tested in future FC harmonization studies.

In this study, we matched individual age and gender across datasets to reduce sample biases. Another covariate of interest in particular to classifying tinnitus from controls is the degree of hearing impairment. Because the condition of tinnitus is highly comorbid with hearing loss, there is a need to dissociate brain connectivity patterns linked to tinnitus from those attributable to hearing loss (Boyen et al., 2014; Husain & Khan, 2023; Schmidt et al., 2013). Our present study is limited by the lack of data for appropriate controls, i.e. those having no tinnitus but only hearing loss. Therefore we focused on classifying individuals having tinnitus but intact hearing, from controls also without hearing impairment. Thus, future harmonization efforts for tinnitus classification should also account for variable hearing thresholds in order to study the condition of tinnitus in the broader population.

## Conclusion

Our findings demonstrate that goal-directed harmonization can effectively integrate resting-state fMRI functional connectivity data across independent datasets while reducing dataset-specific biases and preserving disorder-relevant neural signatures. This approach offers a promising framework for combining existing neuroimaging datasets to investigate altered brain connectivity in tinnitus and other neurological disorders.

## Data availability

HCP-Aging dataset can be accessed by placing a request with the National Institute of Mental Health Data Archive (NDA). Lab dataset on individuals with tinnitus or those classified as controls may be shared upon reasonable request; the corresponding author may be contacted for access.

## Acknowledgement

This work was supported by the Defense Health Agency [Award ID: W81XWH-15-2-0032], the Discovery Partners Institute, the National Center for Supercomputing Applications Illinois Computes program, the Nadine Barrie Smith Memorial Fellowship (granted to S.A.), and the Beckman Graduate Fellowship (granted to S.A.).

## Supplementary Figures

**Supplementary Figure 1:**
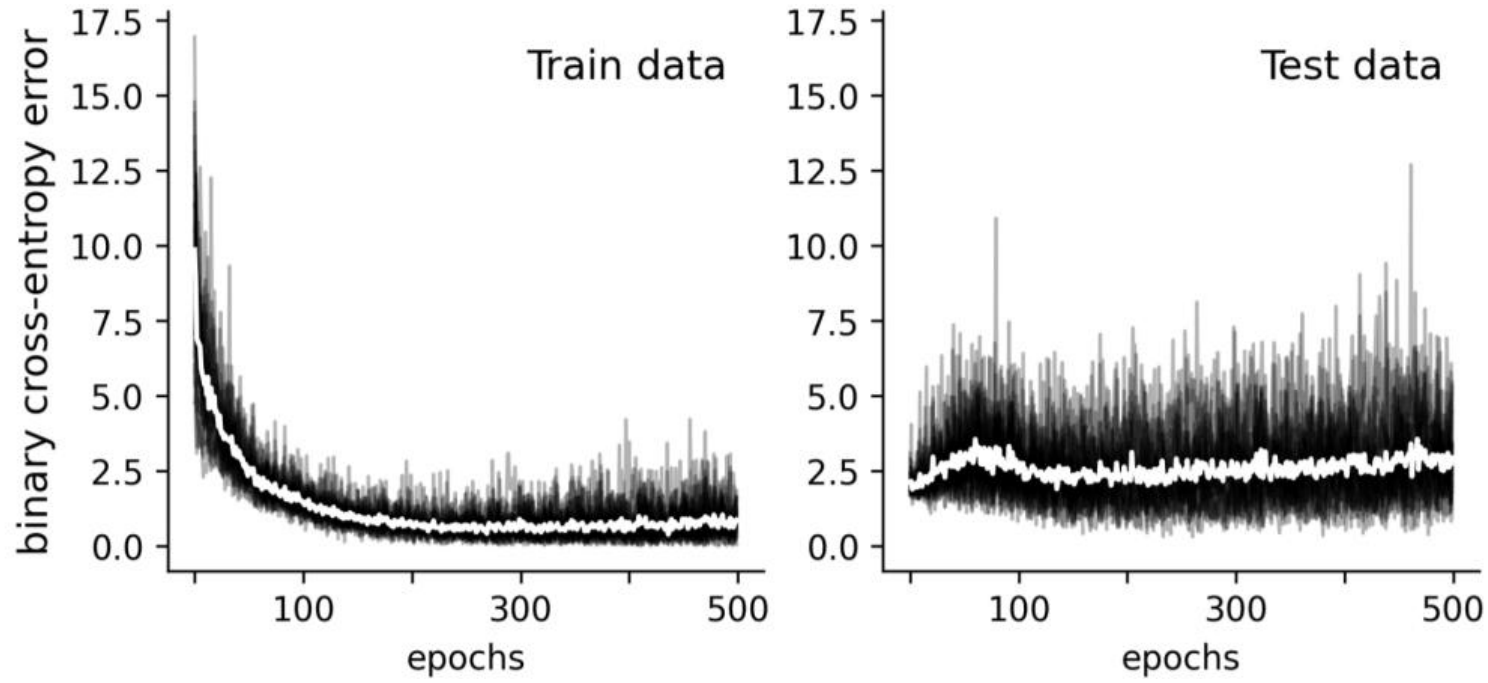
Tinnitus classifier training. Probability prediction error for reconstructed FC across training epochs

**Supplementary Figure 2:**
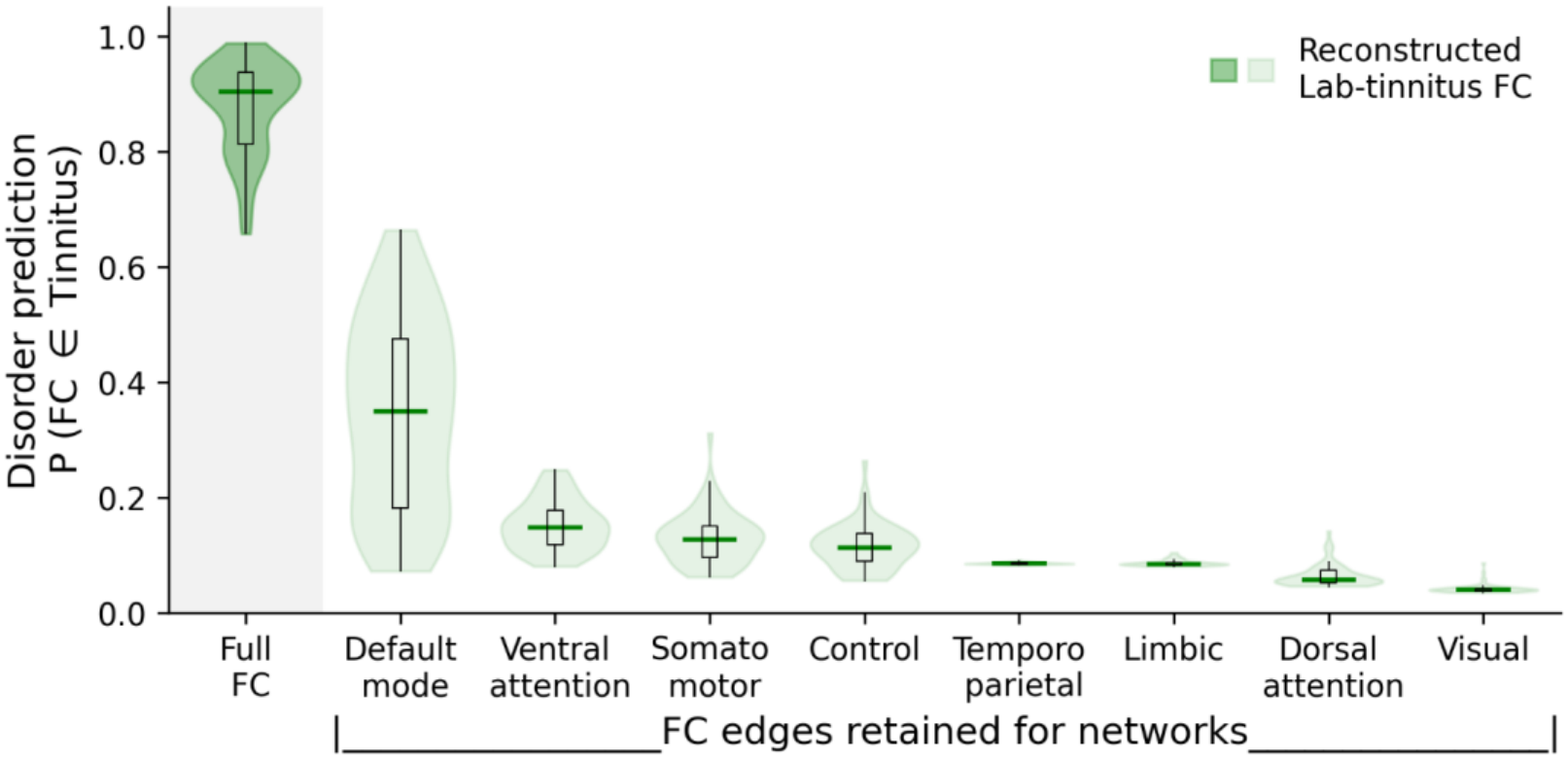
Tinnitus classifier prediction on harmonized Lab tinnitus FC data, when all FC edges are considered (first plot on the left) and when FC edges of ROIs from only one functional network are used (eight plots on the right)

## References

An, L., Chen, J., Chen, P., Zhang, C., He, T., Chen, C., Zhou, J. H., & Yeo, B. T. T. (2022). Goal-specific brain MRI harmonization. NeuroImage, 263, 119570. 10.1016/j.neuroimage.2022.119570

Bookheimer, S. Y., Salat, D. H., Terpstra, M., Ances, B. M., Barch, D. M., Buckner, R. L., Burgess, G. C., Curtiss, S. W., Diaz-Santos, M., Elam, J. S., Fischl, B., Greve, D. N., Hagy, H. A., Harms, M. P., Hatch, O. M., Hedden, T., Hodge, C., Japardi, K. C., Kuhn, T. P., … Yacoub, E. (2019). The Lifespan Human Connectome Project in Aging: An overview. NeuroImage, 185, 335–348. 10.1016/j.neuroimage.2018.10.009

Boyen, K., de Kleine, E., van Dijk, P., & Langers, D. R. M. (2014). Tinnitus-related dissociation between cortical and subcortical neural activity in humans with mild to moderate sensorineural hearing loss. Hearing Research, 312, 48–59. 10.1016/j.heares.2014.03.001

Eckert, M. A., Husain, F. T., M.P. Jayakody D., Schlee, W., & Cederroth, C. R. (2023). An Opportunity for Constructing the Future of Data Sharing in Otolaryngology. Journal of the Association for Research in Otolaryngology, 24(4), 397–399. 10.1007/s10162-023-00908-z

Eggermont, J. J. (2015). Tinnitus and neural plasticity (Tonndorf lecture at XIth International Tinnitus Seminar, Berlin, 2014). Hearing Research, 319, 1–11. 10.1016/j.heares.2014.10.002

Fortin, J.-P., Cullen, N., Sheline, Y. I., Taylor, W. D., Aselcioglu, I., Cook, P. A., Adams, P., Cooper, C., Fava, M., McGrath, P. J., McInnis, M., Phillips, M. L., Trivedi, M. H., Weissman, M. M., & Shinohara, R. T. (2018). Harmonization of cortical thickness measurements across scanners and sites. NeuroImage, 167, 104–120. 10.1016/j.neuroimage.2017.11.024

Fortin, J.-P., Parker, D., Tunç, B., Watanabe, T., Elliott, M. A., Ruparel, K., Roalf, D. R., Satterthwaite, T. D., Gur, R. C., Gur, R. E., Schultz, R. T., Verma, R., & Shinohara, R. T. (2017). Harmonization of multi-site diffusion tensor imaging data. NeuroImage, 161, 149–170. 10.1016/j.neuroimage.2017.08.047

Goodfellow, I. J., Pouget-Abadie, J., Mirza, M., Xu, B., Warde-Farley, D., Ozair, S., Courville, A., & Bengio, Y. (2014). Generative adversarial nets. Advances in Neural Information Processing Systems, 27.

Guntupalli, J. S., Feilong, M., & Haxby, J. V. (2018). A computational model of shared fine-scale structure in the human connectome. PLOS Computational Biology, 14(4), e1006120. 10.1371/journal.pcbi.1006120

Hall, D. A., Fackrell, K., Li, A. B., Thavayogan, R., Smith, S., Kennedy, V., Tinoco, C., Rodrigues, E. D., Campelo, P., Martins, T. D., Lourenço, V. M., Ribeiro, D., & Haider, H. F. (2018). A narrative synthesis of research evidence for tinnitus-related complaints as reported by patients and their significant others. Health and Quality of Life Outcomes, 16(1), 61. 10.1186/s12955-018-0888-9

Husain, F. T., & Khan, R. A. (2023). Review and Perspective on Brain Bases of Tinnitus. Journal of the Association for Research in Otolaryngology, 24(6), 549–562. 10.1007/s10162-023-00914-1

Jarach, C. M., Lugo, A., Scala, M., van den Brandt, P. A., Cederroth, C. R., Odone, A., Garavello, W., Schlee, W., Langguth, B., & Gallus, S. (2022). Global Prevalence and Incidence of Tinnitus. JAMA Neurology, 79(9), 888. 10.1001/jamaneurol.2022.2189

Johnson, W. E., Li, C., & Rabinovic, A. (2007). Adjusting batch effects in microarray expression data using empirical Bayes methods. Biostatistics, 8(1), 118–127. 10.1093/biostatistics/kxj037

Kingma, D. P., & Welling, M. (2019). An Introduction to Variational Autoencoders. Foundations and Trends® in Machine Learning, 12(4), 307–392. 10.1561/2200000056

Langguth, B., Landgrebe, M., Kleinjung, T., Sand, G. P., & Hajak, G. (2011). Tinnitus and depression. The World Journal of Biological Psychiatry, 12(7), 489–500. 10.3109/15622975.2011.575178

Moyer, D., Ver Steeg, G., Tax, C. M. W., & Thompson, P. M. (2020). Scanner invariant representations for diffusion MRI harmonization. Magnetic Resonance in Medicine, 84(4), 2174–2189. 10.1002/mrm.28243

Nastase, S. A., Liu, Y.-F., Hillman, H., Norman, K. A., & Hasson, U. (2020). Leveraging shared connectivity to aggregate heterogeneous datasets into a common response space. NeuroImage, 217, 116865. 10.1016/j.neuroimage.2020.116865

Roberts, L. E., Husain, F. T., & Eggermont, J. J. (2013). Role of attention in the generation and modulation of tinnitus. Neuroscience & Biobehavioral Reviews, 37(8), 1754–1773. 10.1016/j.neubiorev.2013.07.007

Rosemann, S., & Rauschecker, J. P. (2023). Disruptions of default mode network and precuneus connectivity associated with cognitive dysfunctions in tinnitus. Scientific Reports, 13(1), 5746. 10.1038/s41598-023-32599-0

Sabados, A., Kim, C., Rampp, S., Bergherr, E., Buchfelder, M., Schnell, O., & Müller-Voggel, N. (2025). Reducing tinnitus via inhibitory influence of the sensorimotor system on auditory cortical activity. The Journal of Neuroscience, e0581242025. 10.1523/JNEUROSCI.0581-24.2025

Schaefer, A., Kong, R., Gordon, E. M., Laumann, T. O., Zuo, X.-N., Holmes, A. J., Eickhoff, S. B., & Yeo, B. T. T. (2018). Local-Global Parcellation of the Human Cerebral Cortex from Intrinsic Functional Connectivity MRI. Cerebral Cortex, 28(9), 3095–3114. 10.1093/cercor/bhx179

Schmidt, S. A., Akrofi, K., Carpenter-Thompson, J. R., & Husain, F. T. (2013). Default Mode, Dorsal Attention and Auditory Resting State Networks Exhibit Differential Functional Connectivity in Tinnitus and Hearing Loss. PLoS ONE, 8(10), e76488. 10.1371/journal.pone.0076488

Schmidt, S. A., Carpenter-Thompson, J., & Husain, F. T. (2017). Connectivity of precuneus to the default mode and dorsal attention networks: A possible invariant marker of long-term tinnitus. NeuroImage: Clinical, 16, 196–204. 10.1016/j.nicl.2017.07.015

Shahsavarani, S., Schmidt, S. A., Khan, R. A., Tai, Y., & Husain, F. T. (2021). Salience, emotion, and attention: The neural networks underlying tinnitus distress revealed using music and rest. Brain Research, 1755, 147277. 10.1016/j.brainres.2021.147277

Thomas Yeo, B. T., Krienen, F. M., Sepulcre, J., Sabuncu, M. R., Lashkari, D., Hollinshead, M., Roffman, J. L., Smoller, J. W., Zöllei, L., Polimeni, J. R., Fischl, B., Liu, H., & Buckner, R. L. (2011). The organization of the human cerebral cortex estimated by intrinsic functional connectivity. Journal of Neurophysiology, 106(3), 1125–1165. 10.1152/jn.00338.2011

Vanneste, S., van de Heyning, P., & De Ridder, D. (2011). The neural network of phantom sound changes over time: a comparison between recent-onset and chronic tinnitus patients. European Journal of Neuroscience, 34(5), 718–731. 10.1111/j.1460-9568.2011.07793.x

Veterans Benefits Administration. (2023). Annual Benefits Report FISCAL YEAR 2023. https://www.benefits.va.gov/REPORTS/abr/docs/2023-abr.pdf

Watts, E. J., Fackrell, K., Smith, S., Sheldrake, J., Haider, H., & Hoare, D. J. (2018). Why Is Tinnitus a Problem? A Qualitative Analysis of Problems Reported by Tinnitus Patients. Trends in Hearing, 22. 10.1177/2331216518812250

Wineland, A. M., Burton, H., & Piccirillo, J. (2012). Functional Connectivity Networks in Nonbothersome Tinnitus. Otolaryngology–Head and Neck Surgery, 147(5), 900–906. 10.1177/0194599812451414

Yamashita, A., Yahata, N., Itahashi, T., Lisi, G., Yamada, T., Ichikawa, N., Takamura, M., Yoshihara, Y., Kunimatsu, A., Okada, N., Yamagata, H., Matsuo, K., Hashimoto, R., Okada, G., Sakai, Y., Morimoto, J., Narumoto, J., Shimada, Y., Kasai, K., … Imamizu, H. (2019). Harmonization of resting-state functional MRI data across multiple imaging sites via the separation of site differences into sampling bias and measurement bias. PLOS Biology, 17(4), e3000042.. 10.1371/journal.pbio.3000042

